# Imaging Striatal Dopamine Release Using a Non-Genetically Encoded Near-Infrared Fluorescent Catecholamine Nanosensor

**DOI:** 10.1101/356543

**Authors:** Abraham G. Beyene, Kristen Delevich, Jackson Travis Del Bonis-O’Donnell, David J. Piekarski, Wan Chen Lin, A. Wren Thomas, Sarah J. Yang, Polina Kosillo, Darwin Yang, Linda Wilbrecht, Markita P. Landry

## Abstract

Neuromodulation plays a critical role in brain function in both health and disease. New optical tools, and their validation in biological tissues, are needed that can image neuromodulation with high spatial and temporal resolution, which will add an important new dimension of information to neuroscience research. Here, we demonstrate the use of a catecholamine nanosensor with fluorescent emission in the 1000-1300 nm near-infrared window to measure dopamine transmission in *ex vivo* brain slices. These near-infrared catecholamine nanosensors (nIRCats) represent a broader class of nanosensors that can be synthesized from non-covalent conjugation of single wall carbon nanotubes (SWNT) with single strand oligonucleotides. We show that nIRCats can be used to detect catecholamine efflux in brain tissue driven by both electrical stimulation or optogenetic stimulation. Spatial analysis of electrically-evoked signals revealed dynamic regions of interest approximately 2 microns in size in which transients scaled with simulation intensity. Optogenetic stimulation of dopaminergic terminals produced similar transients, whereas optogenetic stimulation of glutamatergic terminals showed no effect on nIRCat signal. Bath application of nomifensine prolonged nIRCat fluorescence signal, consistent with reuptake blockade of dopamine. We further show that the chemically synthetic molecular recognition elements of nIRCats permit measurement of dopamine dynamics in the presence of dopamine receptor agonists and antagonists. These nIRCat nanosensors may be advantageous for future use because i) they do not require virus delivery, gene delivery, or protein expression, ii) their near-infrared fluorescence facilitates imaging in optically scattering brain tissue and is compatible for use in conjunction with other optical neuroscience tool sets, iii) the broad availability of unique near-infrared colors have the potential for simultaneous detection of multiple neurochemical signals, and iv) they are compatible with pharmacology. Together, these data suggest nIRCats and other nanosensors of this class can serve as versatile new optical tools to report dynamics of extracellular neuromodulation in the brain.

## Main

The catecholamines dopamine and norepinephrine are neuromodulators known to play an important role in learning and attention and are implicated in multiple brain disorders.^1–5^ Dopamine, in particular, is thought to play a critical role in learning^6^, motivation^7,8^, and motor control^9^, and aberrations in dopamine neurotransmission are implicated in a wide range of neurological and psychiatric disorders including Parkinson’s disease^10^, schizophrenia^11^, and addiction.^12^

Neuromodulatory neurotransmission is thought to occur on a broader spatial scale than classic neurotransmission, which is largely mediated by synaptic release of the amino acids glutamate and γ-aminobutyric acid (GABA) in the central nervous system. In synaptic glutamatergic and GABAergic neurotransmission, neurotransmitter concentrations briefly rise in the synaptic cleft to mediate local communication between the pre- and postsynaptic neurons through the rapid activation of ligand-gated ion channels.^13^ In contrast, neuromodulators (catecholamines, neuropeptides) may diffuse beyond the synaptic cleft and act via extrasynaptically-expressed metabotropic receptors.^14–19^ Thus, modulatory neurotransmitter activity extends beyond single synaptic partners and enables small numbers of neurons to modulate the activity of broader networks.^20^ The absence of direct change in ionic flux across cell membranes, which is measurable using available tools like electrophysiology or genetically encoded voltage indicators (GEVIs), has necessitated the use of methods borrowed from analytical chemistry such as microdialysis and amperometry to study the dynamics of neuromodulation. However, spatial limitations of fast-scan cyclic voltammetry and spatial and temporal limitations of microdialysis currently make it difficult to interpret how neuromodulators affect the plasticity or function of target neural populations at behaviorally relevant spatiotemporal scales.

Ideally, to understand how neuromodulation sculpts brain activity, we need to develop tools that can optically report modulatory neurotransmitter concentrations in the brain extracellular space (ECS) in a manner that is compatible with pharmacology and other available tools to image neural structure and activity, and minimally interferes with the underlying biology. This goal motivates our effort to design an optical probe that can report extracellular catecholamine dynamics with high spatial and temporal fidelity within a unique near-infrared spectral profile. In this work, we describe the design, characterization, and implementation of a nanoscale near-infrared (nIR) non-genetically encoded fluorescent reporter that allows precise measurement of catecholamine dynamics in brain tissue. This technology makes use of a single wall carbon nanotube (SWNT) non-covalently functionalized with single strand (GT)_6_ oligonucleotides to form the near-infrared catecholamine nanosensor (nIRCat). nIRCats respond to dopamine with *∆F/F* of up to 24-fold in the fluorescence emission window of 1000-1300 nm^48^, a wavelength range that has shown utility for non-invasive through-skull imaging in mice.^21^ First, we show *in vitro* characterization of the nanosensor’s specificity for the catecholamines dopamine and norepinephrine, and demonstrate its insensitivity to GABA, glutamate, and acetylcholine. Second, we use *ex vivo* brain slices containing the dorsal striatum to demonstrate that nIRCats exhibit a fractional change in fluorescence that has the dynamic range and signal-to-noise ratio to report dopamine efflux in response to brief electrical or optogenetic stimulation of dopaminergic terminals. Next, optogenetic stimulation of this preparation is also used to demonstrate the selectivity of the nIRCat nanosensor response to dopaminergic over glutamatergic terminal stimulation. In both stimulation contexts, we show that bath application of dopamine receptor antagonist sulpiride and agonist quinpirole modulates nIRCat signals in a manner consistent with predicted effects of presynaptic D2 autoreceptor manipulation. Finally, we show that the presence of a dopamine reuptake inhibitor yields a prolonged nIRCat fluorescent signal indicating that the sensors report a change in the time course of dopamine diffusion and reuptake in striatal brain tissue. These data indicate that nIRCats provide a unique synthetic tool compatible with pharmacology to interrogate the release, diffusion, and reuptake of neuromodulators in neural tissue.

## Results

### A Near-Infrared Dopamine and Norepinephrine Nanosensor

We report near-infrared fluorescent catecholamine nanosensors (nIRCats) that enable imaging of extrasynaptic catecholamines and their release and re-uptake dynamics in the ECS of brain tissue. Using a previously established nanosensor generation platform^22,23^, synthetic bio-mimetic polymers were pinned onto the surface of intrinsically near-infrared fluorescent single-wall carbon nanotubes (SWNT). The resulting non-covalent nanometer-scale conjugate produced the catecholamine-selective nIRCat. In *in vitro* solution phase experiments (Methods), nIRCats exhibited a maximal change in fluorescence (*∆F/F*) of up to 24 (Figure 1b, 1c) with a dynamic range of 4 orders of magnitude, reporting detectable fluorescence changes from 10 nM to 100 μM dopamine concentration (Figure S1). nIRCats were also sensitive to norepinephrine with a maximal response of *∆F/F*=35 and a similar dynamic range (Figure S1). We found that nIRCats are insensitive to GABA, glutamate, and acetylcholine (Figure 1c) and can report fluctuations in dopamine concentration in the presence of ascorbic acid, which is present in cerebrospinal fluid (Figure S1). Single-molecule imaging revealed that the nIRCat signal in response to repeated perfusions of 10 μM dopamine was reversible upon exposure, an important feature for measuring neuromodulator kinetics (Figure S2). In previous work, we performed stochastic simulations that suggest nIRCats have sufficient sensitivity to detect physiologically relevant fluctuations in dopamine concentration in brain tissue arising from the activity of a single dopaminergic terminal, which can briefly exceed concentrations of 1 μM from the release site in a distance-dependent manner.^24^

**Figure 1.**
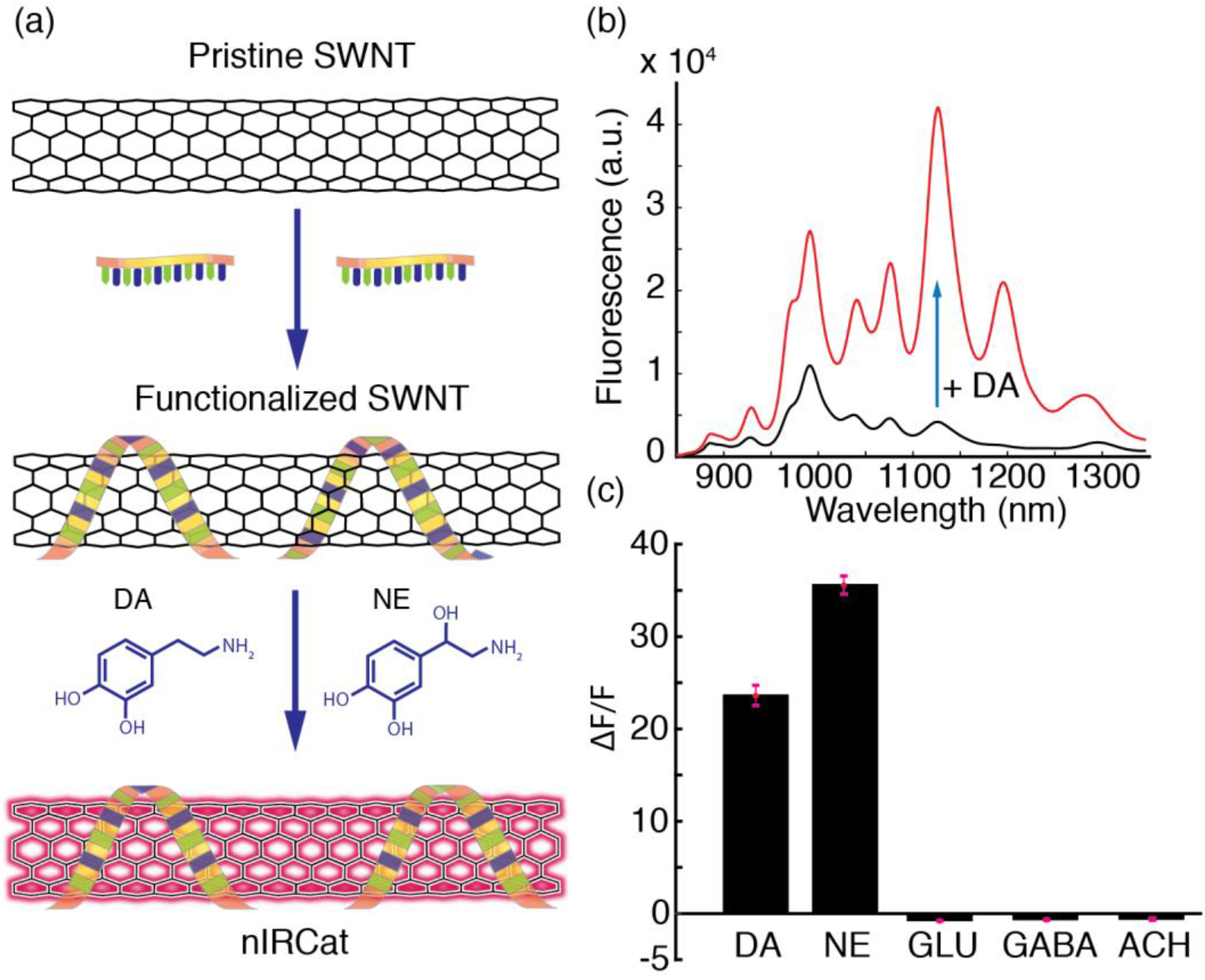
Synthesis and testing of near-infrared catecholamine nanosensors (nIRCats). **(a)** Schematic of optical catecholamine reporters, nIRCats. Pristine SWNT are functionalized with (GT)_6_ oligonucleotides to generate turn-on optical reporters for dopamine (DA) and norepinephrine (NE) **(b)** Fluorescence spectra of nIRCats before (black) and after (red) addition of 10 μM of dopamine in an *in vitro* preparation in phosphate buffered saline (without tissue). Multiple emission peaks correspond to unique SWNT chiralities contained within the multi-chirality mixture. **(c)** Nanosensor optical response to 100 μM dopamine (DA), norepinephrine (NE), glutamate (GLU), γ-aminobutyric acid (GABA), and acetylcholine (ACH) (data from *in vitro* testing). Black bars represent averages from n=3 independent measurements and error bars are calculated as standard deviations of the n=3 measurements.

### Imaging of electrical stimulation-evoked dopamine release in acute striatal brain slices

To determine the efficacy of nIRCats for imaging dopamine in brain tissue, we used brain slices from the dorsal striatum of the mouse. Given that the dorsal striatum is densely innervated by dopaminergic projections from the substantia nigra pars compacta (SNc) but lacks innervation from neurons that release norepinephrine (NE)^25^, we leveraged nIRCats capacity to serve as a dopamine sensor in the striatum. The majority of neurons within the striatum are GABAergic medium spiny neurons (MSNs) with a minority fraction of interneuron populations that include GABAergic and cholinergic interneurons.^26^ Glutamatergic inputs from the cortex and thalamus are the major drivers of MSN activity and dopaminergic terminals in close proximity to these inputs are thought to play an important role in modulating the activity of MSNs and plasticity at striatal synapses.^27^ Due to the composition of local axons, intrastriatal electrical stimulation is predicted to drive the release of a mix of neurotransmitter including GABA, glutamate, acetylcholine, and dopamine, but negligible amounts of other catecholamines like norepinephrine.

Coronal mouse brain slices were prepared as described previously.^28^ Slices were subsequently incubated with 2 mg/L nIRCats for 15 minutes to enable sensors to diffuse into the brain tissue (Figure 2a). Slices were subsequently rinsed to remove excess nIRCats and incubated in standard artificial cerebrospinal fluid (ACSF) for another 15 minutes before imaging. This method, modified from Godin *et al.*^29^ who demonstrated that SWNT localize in the ECS of acute brain slices, enabled even and widespread labeling of the coronal slice, including the dorsal striatum (Figure 2c). Imaging of nIRCat fluorescence modulation in dorsal striatum was accomplished with a custom-built visible and near-infrared microscope to enable concurrent imaging of both visible (400 nm – 750 nm) and near-infrared (750 nm – 1700 nm) wavelengths on the same detector (Figure 2b). Briefly, a 785 nm laser for excitation of nIRCats and mercury bulb for generating brightfield images were directed onto the back focal plane of an epifluorescence upright microscope, and imaging channels were selected using a sliding mirror. Serially, either brightfield or near-infrared images were collected on a Ninox Vis-SWIR 640 broadband camera (Raptor Photonics) with appropriate dichroic filters (Methods) and a 60X water dipping objective (Nikon) providing an imaging field of 178 μm by 142 μm, containing hundreds of dopaminergic terminals.

**Figure 2.**
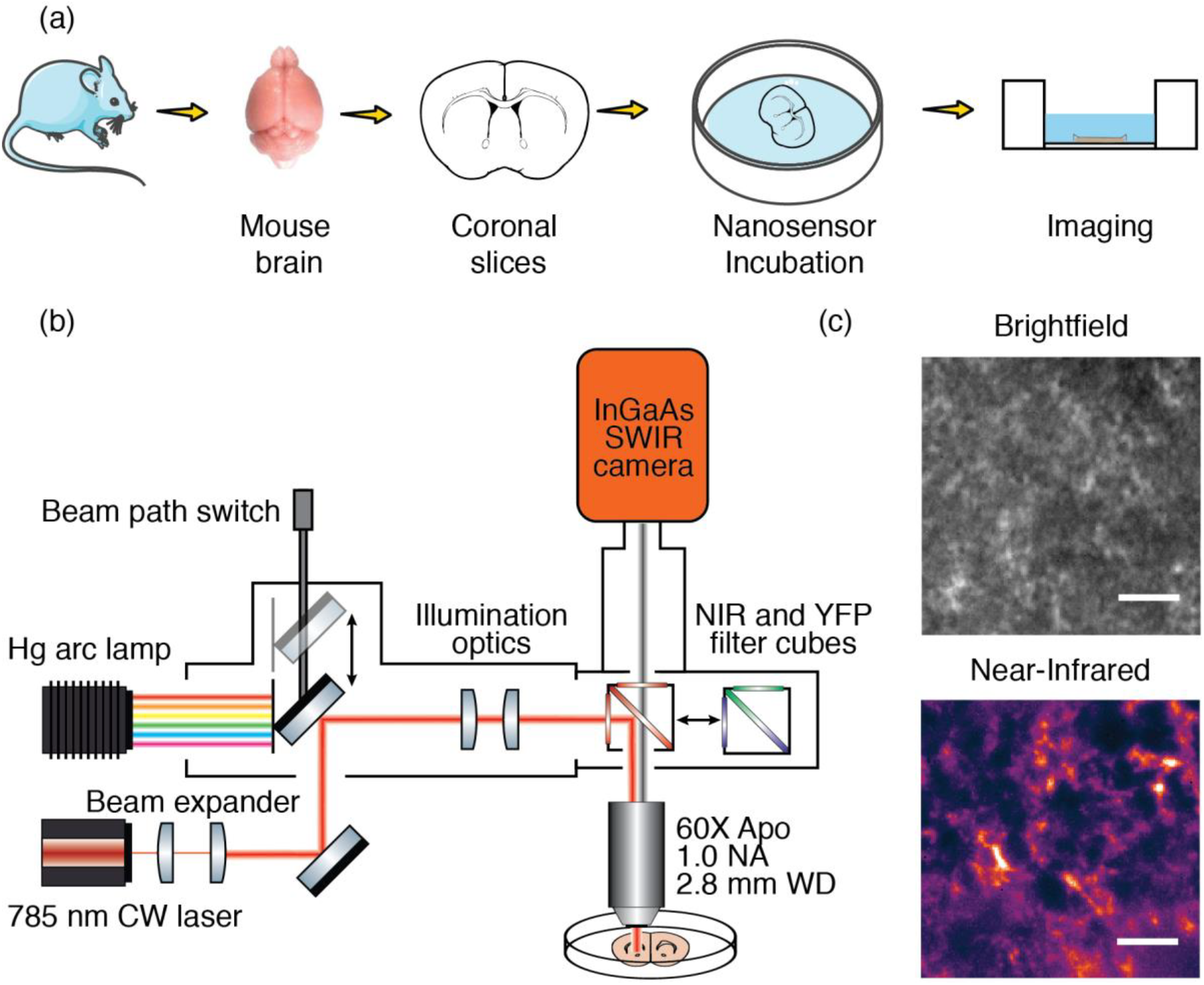
Brain slice nIRCat loading protocol and schematic of visible and near-infrared fluorescence microscopy for imaging nIRCats in acute slices. **(a)** Experimental schematic depicting preparation of acute brain slices and subsequent incubation in 2 mg/L nIRCat solution to load the nanosensors into brain tissue. **(b)** Schematic of visible/near-infrared microscope. A 785 nm CW laser is beam-expanded and co-aligned with a mercury vapor lamp and directed into the objective with dichroic filter cubes. Emitted photons are filtered through a 900 nm long-bass filter and are relayed onto the sensor of a broadband InGaAs camera that is sensitive to both visible and near-infrared wavelengths. **(c)** Dorsomedial striatum from mouse acute slice imaged in brightfield (top) and near-infrared (bottom) after tissue nanosensor loading. Scale bars = 10 μm.

To investigate striatal neuromodulator release with temporal control of tissue stimulation, we used a bipolar stimulating electrode to evoke terminal release within the dorsomedial striatum (stimulus protocol: 3 millisecond wide single square pulses, over 5 biological replicates). We found a single pulse could elicit a nIRCat signal transient, and that increasing the strength of the stimulus led to larger evoked changes in nIRCat *∆F/F* signal, (*∆F/F*)*max* 0.1 mA = 0.047 ± 0.025; 0.3 mA = 0.122 ± 0.026; and 0.5 mA = 0.2 ± 0.033; mean ± s. d., n=5 for all measurements, p=0.008 between 0.1 mA vs. 0.3 mA, p=0.008 between 0.3 mA vs. 0.5 mA) (Figure 3b).

**Figure 3.**
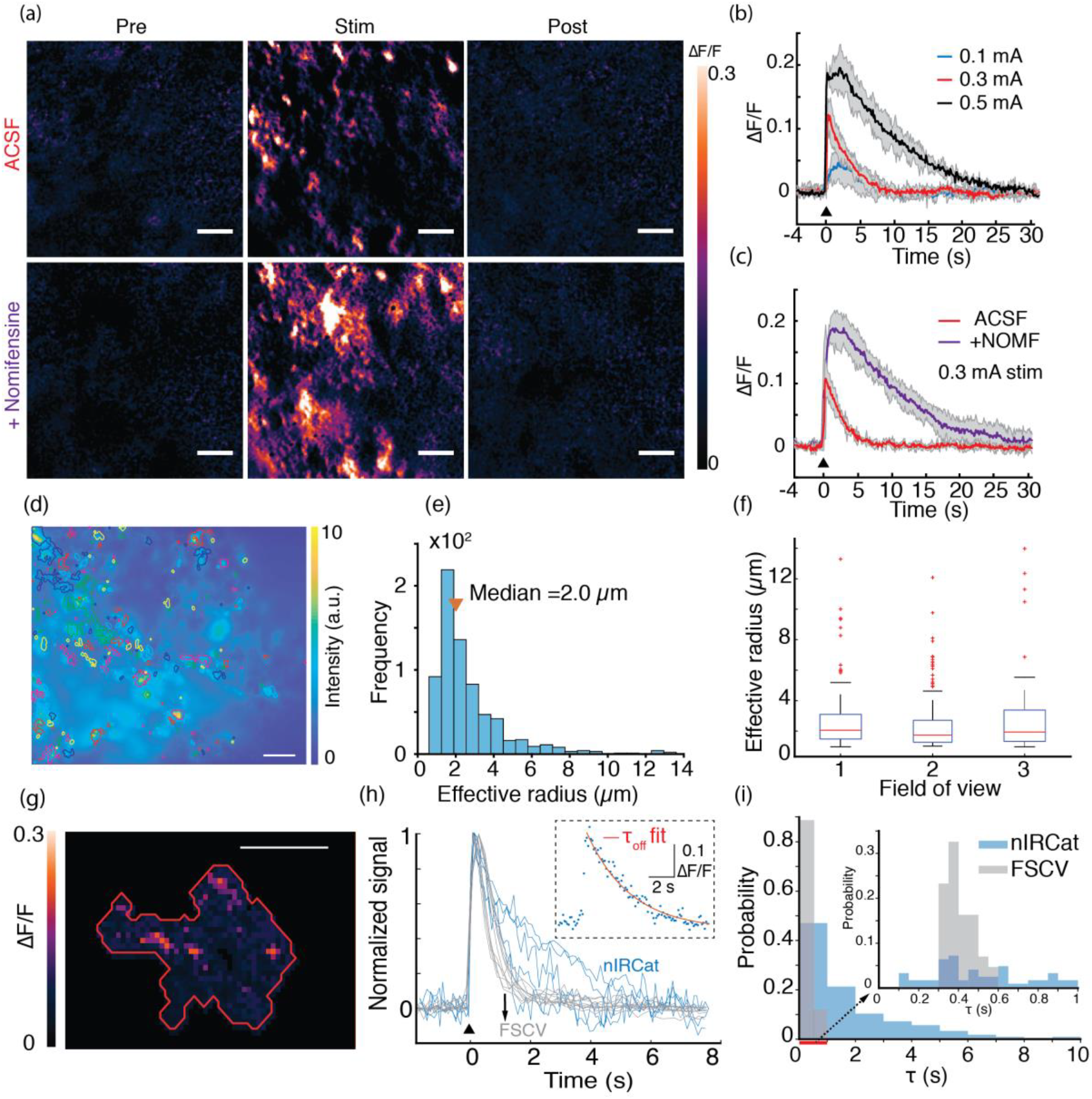
Imaging and spatiotemporal analysis of dopamine release evoked by electrical stimulation in acute striatal slices. **(a)** Nanosensor *∆F/F* imaging of the same acute brain slice field of view upon electrical stimulation by 0.3 mA in standard ACSF (top row) and in ACSF plus 10 μM nomifensine (bottom, +Nomifensine). Three still frames are presented from each movie. “Pre” is a representative frame before electrical stimulation is applied, “Stim” represents frame corresponding to peak *∆F/F* following stimulation, and “Post” is a representative frame after nIRCat fluorescence has returned to baseline. Scale bars = 10 μm. **(b)** Nanosensor fluorescence modulation scales with single pulse electrical stimulation amplitudes. Field of view mean traces and standard deviation bands are presented for three stimulation amplitudes of 0.1 mA, 0.3 mA, and 0.5 mA (n=5) **(c)** Time traces of *∆F/F* for 0.3 mA single pulse stimulation in standard ACSF (red) and in ACSF plus 10 μM nomifensine (purple, +NOMF). Mean traces with standard deviation bands are presented (n=3). **(d)** A single frame from a time series of imaging in the dorsomedial striatum showing the entire field of view, overlaid with ROIs identified from an automated program from per-pixel *∆F/F* stack projections of nIRCat fluorescence modulation (see Methods). Color bar represents nIRCat fluorescence intensity of labeled tissue. Scale bar = 20 μm. **(e)** Frequency histogram of ROI sizes depicted in (d), exhibiting a log-normal distribution with median ROI size of 2 μm. **(f)** ROI size distribution from three different fields of view (representing n = 3 biological replicates) in the dorsal striatum, each stimulated separately by 0.3 mA stimulation. In each of these fields of view, ROIs showed similar median size and size distribution even when compared across biological replicates. Box plot definitions: red-line = median, edges of box: 25^th^ and 75^th^ percentiles, top and bottom lines: minimum and maximum values of non-outlier data, red points: outlier data **(g)** A higher magnification view of an ROI with an effective radius of 5 μm. Maximum *∆F/F* projection of the ROI shows presence of smaller fluorescence hotspots within the ROI. Scale bar = 5 μm. **(h)** Overlay of representative normalized FSCV (gray) and nIRCat (blue) traces showing that nIRCat ROI signals exhibit a wider diversity of decay kinetics. Inset: An example of nIRCat experimental data (blue dots) fitted to first order kinetics (red line) to compute decay time constants (τ). **(i)** Normalized frequency histogram of τ’s computed from FSCV and nIRCat individual ROI time traces. Data from n= 4 fields of view representing n=2 biological replicates is pooled. Medians of each distribution are: nIRCats τ = 1.1 s and FSCV τ = 0.4 s.

To further test if evoked nIRCat signals tracked striatal dopamine release and reuptake kinetics, we investigated the effect of nomifensine, a dopamine reuptake inhibitor that slows the clearance of dopamine from the ECS by competitively binding to dopamine transporters (DATs). Addition of 10 μM nomifensine to the bath yielded nIRCat signal with higher peak fluorescence modulation (*(∆F/F*)_*max*_ = 0.108 ± 0.029 vs. 0.189 ± 0.023; mean ± s. d., n=3, p=0.0178) and a prolonged fluorescent signal compared to signals obtained in ACSF from the same field of view (decay time constant, τ=2.43 ± 0.24 s vs. 10.95 ± 1.15 s; mean ± s. d., n=3, p=0.0002) (Figure 3a top vs. bottom row, Figure 3c).

To identify nIRCat fluorescence change hotspots (i.e., regions of high ∆F/F), we analyzed our video-rate acquisitions using a custom-built program that accounted for background fluorescence and identified regions with fluctuations in fluorescence intensity in the post-stimulation epoch (see Methods for details). We defined nIRCat ∆F/F hotspots as regions of interest (ROIs) based on a per-pixel stack projection of maximal ∆F/F in the imaging time series. The ROIs identified were found to vary in size from 1 μm to 15 μm (Figure 3d, 3e). We found that ∆F/F hotspots do not necessarily correspond to high nIRCat labeling of the brain tissue, suggesting that the hotspot is a consequence of variation in dopamine release and not nanosensor loading in the tissue (Figure 3d, Figure S3 a-c). Using data from single pulse electrical stimulation experiments, the program identified ROIs whose size distribution was observed to exhibit a log-normal distribution with median ROI size of 2μm (Figure 3e). Repeat stimulations with the same stimulation amplitude in fields of view of the dorsomedial striatum across biological replicates generated similar size distributions (Figure 3f). Closer examination of several larger ROIs (> 5μm) suggested these may be comprised of smaller hot-spots in close proximity (Figure 3g, Figure S3 d-e).

For further examination of the temporal resolution of nIRCats, we compared the temporal profile of evoked transients measured with nIRCats to transients measured with fast scan cyclic voltammetry (FSCV). FSCV is a technique that has been widely used to measure temporal catecholamine dynamics both *in vivo* and *in vitro* in the striatum and other brain areas.^30–32^ FSCV and nIRCat experiments were conducted on separate experimental rigs with the same solutions, temperature settings, electrodes, and stimulation parameters. Evoked transients measured with FSCV and nIRCat fluorescence emission showed comparable temporal profiles in the rising phase (latency to peak: FSCV = 0.25 ± 0.0 s vs. nIRCat = 0.40 ± 0.18 s; mean ± s. d., n=4 fields of view from 2 biological replicates, p=0.23). Meanwhile, nIRCat signals exhibited a wider diversity of decay kinetics (τ: FSCV = 0.51 ± 0.08 s vs. nIRCats = 2.43 ± 0.24 s; mean ± s. d. n=4 fields of view from 2 biological replicates, p=0.0002). A subset of ROIs exhibited decay time constants that overlapped with, or were faster than, those of FSCV signals (Figure 3h, 3i).

We next evaluated the ability of nIRCats to detect dopamine in the presence of dopamine receptor (DR) drugs. In *in vitro* solution phase experiments (without biological tissue), we found that nIRCat fluorescence intensity was not modulated by exposure to 1 μM concentration of DRD2 antagonists sulpiride, haloperidol, the DRD2 agonist quinpirole, or the DRD1 antagonist SCH-23390 (Figure 4a). Furthermore, *in vitro* solution phase dopamine-induced nIRCat fluorescence signals were not altered in the presence of these same dopamine receptor drugs (Figure 4a).

**Figure 4.**
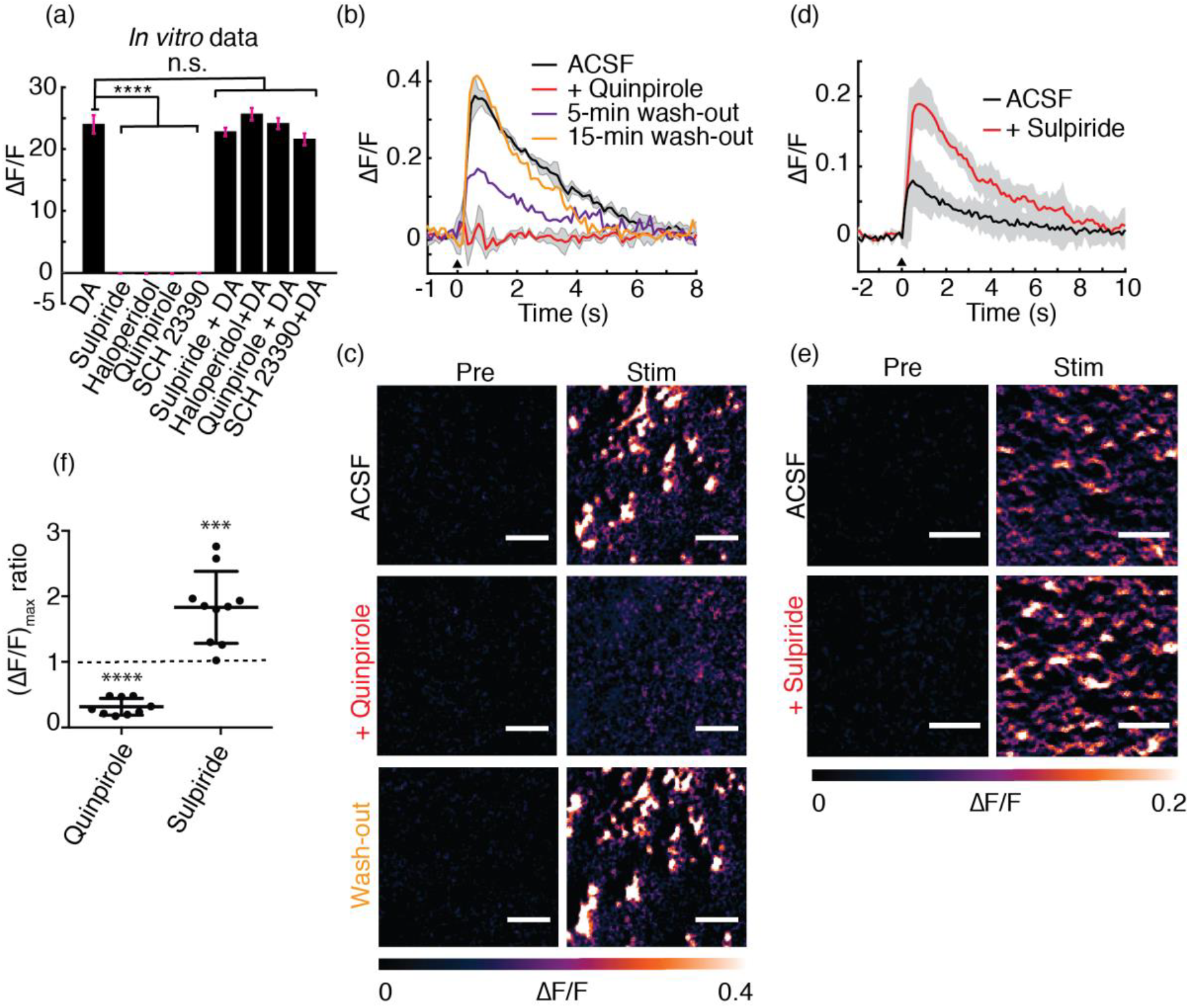
Imaging dopamine release in the presence of dopamine receptor drugs. **(a)** *In vitro* solution phase maximal *∆F/F* of nIRCat in presence of 100 μM dopamine (DA), the dopamine receptor drugs sulpiride, haloperidol (DRD2 antagonists), quinpirole (DRD2 agonist), and SCH 23390 (DRD1 antagonist), and dopamine receptor drugs + DA. Addition of 1 μM drug quantities did not induce nIRCat fluorescence modulation in the absence of DA (p < 0.0001 compared to DA *∆F/F*). Subsequent addition of DA to drug-incubated nIRCat solutions produced *∆F/F* responses indistinguishable from DA-only responses. Error bars represent standard deviations from n=3 measurements **(b, c)** In slice, quinpirole (1 μM) abolished nIRCat fluorescence modulation in response to a single electrical pulse (0.5 mA) (red trace) compared to pre-drug ACSF (black trace) but recovered following drug wash-out (purple and orange traces). **(d,e)** Sulpiride (1 μM) enhanced nIRCat fluorescence modulation in response to single electrical pulse stimulation, yielding brighter nIRCat ∆F/F hotspots compared to drug-free ACSF. **(f)** Quinpirole depressed nIRCat fluorescence modulation (p<0.0001), whereas sulpiride facilitated it (p=0.001) in n=3 biological replicates. Individual data points represent *(∆F/F)_max_* ratio of the average trace collected in same field of view (post-drug application/pre-drug application). Scale bars = 10 μm. All error bands represent standard deviation from the mean trace.

In *ex vivo* brain slices (dorsal striatum), application of quinpirole (1 μM) abolished nIRCat fluorescent transients in response to single electrical pulse stimulation which was rescued following 15-minute drug wash-out (Figure 4b, 4c). Conversely, application of sulpiride (1μM) significantly increased nIRCat ∆F/F (Figure 4d, 4e). The effects of quinpirole and sulpiride were reproducible in n=3 biological replicates (Figure 4f). Importantly, the effects of D2 receptor pharmacology on nIRCat fluorescence modulation by dopamine were absent *in vitro* and only present in brain tissue. In brain tissue, presynaptic dopamine receptors play a critical role in regulating dopamine release, and our results are consistent with powerful inhibition of dopamine efflux by the DRD2 agonist quinpirole and facilitation of dopamine efflux by the DRD2 antagonist sulpiride.

### Imaging of optogenetically-evoked dopamine release in acute striatal brain slices

To further confirm striatal nIRCat nanosenor signals were indeed reporting dopamine release, we compared channelrhodopsin (ChR2) stimulation of cortical glutamatergic and nigrostriatal dopaminergic terminals in the dorsal striatum. Acute striatal brain slices were prepared from mice virally transfected to express the light sensitive cation channel ChR2 in either dopaminergic terminals (targeted by viral injection in the midbrain in DAT-cre mice; ChR2-DA) (Figure 5c, Figure S4) or glutamatergic terminals of the striatum (targeted by viral injection in the frontal cortices, ChR2-GLU) (Figure 5a). Upon optical stimulation of ChR2-DA terminals with a 473 nm laser (5 pulses at 25 Hz, 1 mW/mm^2^) in the dorsal striatum, we observed significant fluorescence modulation of nIRCat signal (Figure 5d). When optogenetic stimulation instead targeted glutamatergic terminals, fluorescent nIRCat signals did not rise above baseline fluctuation, although we could observe light-evoked excitatory postsynaptic currents (EPSCs) in striatal MSNs (Figure 5b).

**Figure 5.**
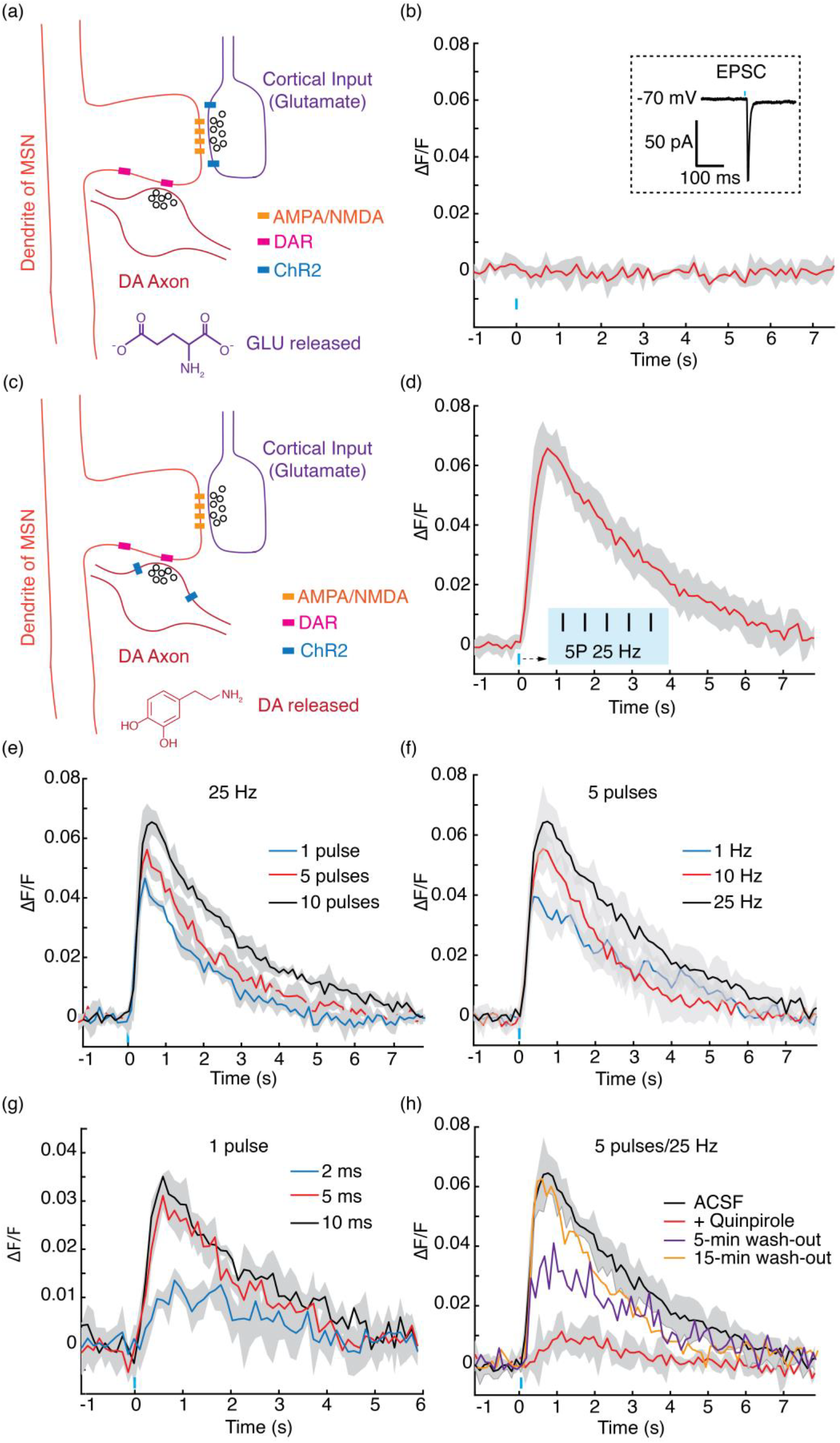
Optogenetic release of dopamine imaged by nIRCats. **(a)** Schematic of ChR2 expression in cortical glutamatergic terminals of the dorsal striatum. **(b)** No nIRCat fluorescence modulation was observed after stimulation of glutamate terminals. Inset: glutamate release was confirmed by excitatory postsynaptic current on medium spiny neuron. **(c)** Schematic of ChR2 expression in nigrostriatal dopaminergic terminals of the dorsal striatum. **(d)** Stimulation of dopaminergic terminals resulted in nIRCat fluorescence modulation. Stimulation protocol in (b) and (d) was 5 pulses at 25 Hz and power flux of 1 mW/mm^2^ and each pulse had a duration of 5ms. (e). nIRCat ∆F/F in response to increasing number of pulses delivered at 25 Hz, 5 ms pulse duration. (f) nIRCat ∆F/F in response to increasing pulse frequency (1, 10, 25 Hz) of 5 pulses. Each pulse had a duration of 5 ms. (g) nIRCat ∆F/F in response to single pulses of 2 ms, 5 ms and 10 ms duration. (h) Bath application of 1 μM of quinpirole suppresses dopamine release and results in depressed nIRCat ∆F/F. Drug wash-out rescues dopamine release and nIRCat ∆F/F. All error bands represent standard deviation from the mean trace.

Returning to ChR2-DA stimulation, we next varied the number of stimulation pulses (5 ms pulse duration, 25 Hz, 1 mW/mm^2^) and observed scaling in nIRCat ∆F/F amplitude from 1 pulse to 10 pulses (p=0.005), and trend level differences between 1 pulse and 5 pulses (p=0.0645) and 5 pulses and 10 pulses (p=0.086) (Figure 5e). When we varied the pulse frequency while holding the number of pulses constant at 5, we observed scaling with significant differences detectable between 1 Hz and 10 Hz (p=0.036). The amplitude difference between 10 Hz and 25 Hz did not reach significance (p=0.179) (Figure 5f). In single pulse experiments in which we varied pulse width, nIRCat fluorescence responses scaled with pulse duration. This effect was significant when comparing 2 ms to 5 ms (p=0.002) but the difference from 5 ms to 10 ms did not reach significance (p=0.055) (Figure 5g).

Finally we tested the effect of dopaminergic pharmacological agents on optogenetically evoked dopamine release (ChR2-DA). Bath application of quinpirole (1 μM) powerfully suppressed nIRCat fluorescence (p<0.0001), and this effect was reversed after drug washout (Figure 5h). Nomifensine (10 μM) enhanced nIRCat signal decay time, consistent with the predicted slowing of dopamine clearance from the ECS (Figure S5).

## Conclusion

In order to understand how neuromodulation alters the plasticity and activity of distinct populations of neurons, there is need for new optical tools that can measure the extracellular dynamics of neuromodulator release and reuptake at spatiotemporal resolution commensurate with methods used to record neural activity (e.g. electrophysiology and calcium imaging). Here, we demonstrated the feasibility of using of a non-genetically encoded fluorescent sensor, nIRCat, which enables optical detection of catecholamine release and reuptake with sub-second temporal and with micrometer spatial resolution. We used electrical and optogenetic methods in *ex vivo* brain slices to demonstrate that nIRCat fluorescent signals can successfully report differences in evoked dopamine release and pharmacologically induced changes in dopamine dynamics in striatal brain tissue.

Here, we focused nIRCat imaging experiments within the dorsal striatum, a region that receives dense dopaminergic innervation and negligible norepinephrine innervation.^25^ Therefore, while nIRCats are not selective for dopamine over norepinephrine, nIRCats effectively function as a dopamine sensor within the context of the striatum. Given that striatal dopamine regulates fundamental processes including motor function, motivation and learning, nIRCats represent an important addition to the neuroscience investigative toolkit.

While other tools are emerging to optically report dopamine fluctuations via cell-surface engineered proteins,^33,34^ nIRCats are likely to fulfill a niche amongst currently available methods for detecting dopamine neurotransmission due to their unique near-infrared fluorescence, the fact that they do not rely on genetic delivery and expression, and the functionality of nIRCats in the presence of pharmacological dopamine receptor ligands. This is in contrast to receptor-based fluorescent sensors which currently cannot report on endogenous dopamine dynamics in the presence of ligands to the engineered receptor.^33,34^ Furthermore, the synthetic nature of nIRCats affords complete elimination of potential GPCR-mediated residual signaling that may be present in protein-based optical probes. nIRCats also offer spatial advantages over FSCV, and our initial experiments suggest that the temporal resolution of the nIRCat signal can be comparable to that of FSCV. nIRCat decay profiles exhibit a wider range than that observed from FSCV data and included a significant number of ROIs that showed seconds-long time constants. It is debatable if these results capture the unique spatial properties of specific striatal ROIs or are related to tool differences alone. nIRCats, unlike FSCV probes, should sample catecholamine concentration at a single point in space, such that each distributed nIRCat construct can act as a probe within the ECS and therefore yield a “higher resolution” picture of catecholamine dynamics. Future experiments will investigate how the heterogeneity of nIRCat signals (amplitude and kinetics) relates to structural and functional properties at dopamine terminals and within the ECS. We predict that new optical tools for measuring dopamine dynamics with high spatial resolution will be enable new insights into the regulation of dopamine release and reuptake at the level of individual synapses.^35–37^

In the future, we see potential for expansion of a larger family of SWNT based near-infrared nanosensors (nIRNS) similar to nIRCats for multiple neurochemical imaging applications. Several lines of evidence illustrate their future potential. First, nIRNS are easily functionalized with a wide range of synthetic molecular recognition moieties, affording fine control of their surface functional elements and their interactions with the local chemical environment.^38–40^ SWNT fluorescence can be finely tuned to monochromatic emission in the near-infrared II (1000-1700 nm) window by controlling the SWNT chirality.^41^ This chirality-dependent fluorescence in the near-infrared II window provides further avenues for designing color-specific responses to multiple molecular analytes simultaneously, thereby affording synthesis of ratiometric and multiplexed analyte imaging platforms, as we have shown previously.^42^ Second, SWNT-based nanosensors rely on near-infrared fluorescence, which greatly reduces the impact of tissue scattering in the emission window and therefore may enable through-cranium imaging.^21^ nIRNS are compatible with multi-photon imaging with 1600 nm excitation^43^ and as such could permit nanoscale imaging of intact neuronal structures pending parallel developments in all-infrared microscopy, as has been shown with visible wavelength-emitting fluorophores.^44^ Third, nIRNS have robust photostability allowing for use in long-term imaging experiments.^45^ Fourth, because nIRNS are not genetically-encoded, they could enable use in species where gene delivery and protein expression is too difficult, time consuming, or undesirable. Finally, nanosecond-scale binding kinetics and the nanoscale dimensions of nIRNS^48^, are likely to enable generation of nanosensors with improved temporal and spatial resolution. In sum, nIRCats are versatile catecholamine probes amenable to multiplexing with existing tools for concurrent investigation of dopaminergic neuromodulation with other core mechanisms of brain function.

## Methods

### Nanosensor synthesis

(GT)_6_ oligonucleotides were purchased from Integrated DNA Technologies (IDT, Standard Desalting). HiPCo SWNT were purchased from NanoIntegris (Batch # HR27-104). (GT)_6_- SWNT colloidal suspension (nIRCat) was prepared by mixing 1 mg of (GT)_6_ and 1 mg of SWNT in 1 mL of a 100 mM NaCl solution. The solution was bath sonicated (Branson Ultrasonic 1800) for 10 minutes and probe-tip sonicated for 10 minutes at 5 W power (Cole Parmer Ultrasonic Processor, 3 mm tip diameter) in an ice-bath. The sonicated solution was incubated at room temperature for 30 minutes. The product was subsequently centrifuged at 16,000 g (Eppendorf 5418) for 90 minutes to remove unsuspended SWNT bundles and amorphous carbon, and the supernatant was recovered for characterization and use. Each nanosensor suspension was stored at 4°C until use.

### Nanosensor characterization

To characterize nIRCats post-synthesis, the full visible and near-infrared absorption spectrum was taken for each nanosensor batch (UV-VIS-NIR spectrophotometer, Shimadzu UV-3600 Plus) or UV-VIS (ThermoFisher Scientific Genesys 20). SWNT concentrations of as-made nanosensor batches were determined using absorbance at 632 nm (UV-VIS) with an extinction coefficient of ε = 0.036 (mg/L)^−1^ cm^−1^. Full spectrum absorbance measurements were made with UV-VIS-NIR after dilution to 5 mg/L SWNT concentration in 100 mM NaCl. For fluorescence spectroscopy, each sensor batch was diluted to a working concentration of 5 mg/L in 100 mM NaCl, and aliquots of 198 μL were placed in each well of a 96-well plate (Corning). Fluorescence measurements were obtained with a 20 X objective on an inverted Zeiss microscope (Axio Observer.D1) coupled to a Princeton Instruments spectrograph (SCT 320) and liquid nitrogen cooled Princeton Instruments InGaAs linear array detector (PyLoN-IR). A 721nm laser (OptoEngine LLC) was used as the excitation light source for all characterization experiments.

### Neurotransmitter and dopamine receptor drug screening

For neurotransmitter response screens, we collected the near-infrared fluorescence spectrum from 198 μL aliquots of nanosensor (5 mg/L SWNT concentration) before and after addition of 2 μL of 10 mM solutions of each analyte neurotransmitter (for a 100 μM final analyte concentration in each well of a 96-well plate). All neurotransmitter analytes were purchased from Sigma-Aldrich. Neurotransmitter analytes were incubated for 5 minutes before taking post-analyte fluorescence measurements. Responses were calculated for the integrated fluorescence count as *∆F/F_0_* = *(F-F_0_)/F_0_*, where *F_0_* is total fluorescence before analyte addition and *F* is total fluorescence after analyte addition or for peak fluorescence change corresponding to the (9,4) SWNT chirality (~1128 nm center wavelength). All measurements were made in triplicate. Reported results are mean and standard deviations of the triplicate measurements. All nIRCat nanosensor batches were tested for catecholamine responses prior to use for tissue catecholamine imaging. Dopamine receptor drugs were purchased from Tocris (quinpirole and sulpiride), abcam (SCH 23390) and Sigma-Aldrich (haloperidol). nIRCat fluorescence modulation to dopamine receptor drugs were measured after addition of 1 μM drug quantities (final concentration in well) in each well. Post-drug fluorescence spectra were taken after 5- minute drug incubation. To measure nIRCat response to dopamine in the presence of drugs, dopamine aliquots were added to each drug-incubated well to obtain 100 μM dopamine, and post-dopamine fluorescence spectra were taken after an additional 5-minute incubation period.

### Nanosensor reversibility testing

A #1.5 glass coverslip was functionalized with (3-Aminopropyl) triethoxysilane (APTES, Sigma Aldrich) by soaking in 10% APTES in ethanol for 5 min. The coverslip was then rinsed with DI water and left to dry. The coverslip was then fixed onto an ibidi sticky-Slide VI 0.4 forming 6 microfluidic channels. First, 100 μL of PBS was pipetted through a channel. Next, the channel was filled 50 μL of a 5 mg/L solution of nIRCats and left to incubate at room temperature for 5 min. The channel was rinsed using three 50 μL PBS washes, keeping the channel filled with solution at all times. The surface immobilized nIRCats in PBS were imaged on an epifluorescence microscope with 721 nm excitation and a Ninox VIS-SWIR 640 camera (Raptor). One end of the flow channel was connected to a syringe pump (Harvard Appartus) using Luer lock fittings. Prior to the start of image acquisition, the opposite flow reservoir was filled with PBS and the pump was set to refill mode at a volumetric flow rate of 40 μL min^−1^. Once the liquid in the reservoir was depleted, 40 μL of 10 μM dopamine in PBS was added. The process was repeated using alternating additions of 80 μL of PBS washes and 40 μL of dopamine solution.

### Acute slice preparation and nanosensor labeling

Mice were C57 Bl/6 strain, 60 days old, and both male and female mice were used. Mice were group housed after weaning at P21 and kept with nesting material on a 12:12 light cycle. All animal procedures were approved by the UC Berkeley Animal Care and Use Committee (ACUC). Acute brain slices were prepared using established protocols.^28^ Briefly, mice were deeply anesthetized via intraperitoneal injection of ketamine/xylazine cocktail and transcardial perfusion was performed using ice-cold cutting buffer (119 mM NaCl, 26.2 mM NaHCO_3_, 2.5 mM KCl, 1mM NaH_2_PO_4_, 3.5 mM MgCl_2_, 10 mM Glucose, 0 mM CaCl_2_), after which the brain was rapidly extracted. The cerebellum and other connective tissues were trimmed using a razor blade and the brain was mounted onto the cutting stage of a vibratome (Leica VT 1200S). Coronal slices (300 μm in thickness) including the dorsal striatum were prepared. Slices were incubated at 37°C for 60 minutes in oxygen saturated ACSF (119 mM NaCl, 26.2 mM NaHCO_3_, 2.5 mM KCl, 1mM NaH_2_PO_4_, 1.3 mM MgCl_2_, 10 mM Glucose, 2 mM CaCl_2_) before use. Slices were then transferred to room temperature for 30 minutes before starting imaging experiments and were maintained at room temperature for the remainder of experimentation.

For nanosensor labeling, slices were transferred into a small volume brain slice incubation chamber (Scientific Systems Design, Inc., AutoMate Scientific) and kept under oxygen saturated ACSF (total 5 mL volume). 100 μL of 100 mg/L nIRCat nanosensor was added to the 5mL volume and the slice was incubated in this solution for 15 minutes. The slice was subsequently recovered and rinsed in oxygen saturated ACSF to wash off nIRCats that did not localize into the brain tissue. The rinsing step was performed by transferring the slice through 3 wells of a 24 well plate (5 seconds in each well) followed by transfer to the recording chamber with ACSF perfusion for a 15-minute equilibration period before starting the imaging experimentation. All imaging experiments were performed at 32°C.

### Acute slice preparation for FSCV recording

Acute slices were prepared as described previously. Extracellular dopamine concentration evoked by local electrical stimulation was monitored with FSCV at carbon-fiber microelectrodes (CFMs) using Millar voltammeter. CFMs were ~ 7 μm in diameter encased in glass capillary pulled to form a seal with the fiber and cut to final tip length of 70-120 μm. The CFM was positioned ~100 μm below the tissue surface at a 45-degree angle. A triangular waveform was applied to the CFM scanning from −0.7 V to +1.3 V and back, against Ag/AgCl reference electrode at a rate of 800 V/s. Evoked dopamine transients were sampled at 8 Hz, and data were acquired at 50 kHz using AxoScope 10.5 (Molecular Devices). Oxidation currents evoked by electrical stimulation were converted to dopamine concentration from post-experimental calibrations. Recorded FSCV signals were identified as dopamine by comparing oxidation (+0.6 V) and reduction (−0.2 V) potential peaks from experimental voltammograms with currents recorded during calibration with 2 μM dopamine dissolved in ACSF. For stimulation, a bipolar stimulation electrode (FHC CBAEC75) was positioned on top of the brain slice and approximately 100 μm away from the CFM. Following 30-minute slice equilibration in the recording chamber, dopamine release was evoked using a square pulse (0.3 mA pulse amplitude, 3 ms pulse duration) controlled by Isoflex stimulus isolator (A.M.P.I) and delivered out of phase with the voltammetric scans. Stimulation was repeated 3 times. To compare FSCV and nIRcat data, each signal was normalized against its peak value ([DA]_*max*_ or (∆F/F)_*max*_) and co-aligned at stimulation time. Latency to peak were computed as *t_peak_* - *t_stim_* where *t_peak_* is the time at which peak signal is attained and *t_stim_* is time of stimulation. Decay time constants (τ) were computed from model fits to a first order decay process.

### Microscope construction and slice imaging

*Ex vivo* slice imaging was performed with a modified upright epifluorescent microscope (Olympus, Sutter Instruments) mounted onto a motorized stage. Nanosensor excitation was supplied by a 785 nm, CW, DPSS laser with adjustable output power to a maximum of 300 mW and a near TEM00, top hat beam profile (OptoEngine LLC). The beam was expanded using a Keplerian beam expander comprised of two plano-convex lenses (Thorlabs, *f*=25 mm, *f*=75 mm, AR coating B) to a final beam diameter of approximately 1 cm. The beam was passed through a custom fluorescence filter cube (excitation: 800 nm shortpass (FESH0800), dichroic: 900 longpass (DMLP990R), emission: 900 longpass (FELH0900), Thorlabs) to a 60X Apo objective (1.0 NA, 2.8 mm WD, water dipping, high NIR transmission, Nikon CFI Apo 60XW NIR). Emission photons collected from the sample were passed through the filter cube and were focused onto a two-dimensional InGaAs array detector (500-600 nm: 40% quantum efficiency (QE); 1000-1500 nm: >85% QE; Ninox 640, Raptor Photonics) and recorded using Micro-Manager Open Source Microscopy Software.46 Laser power was adjusted to maximize collected photons and fill the pixel bit depth on the detector but did not exceed 70 mW at the objective back focal plane. YFP fluorescence was imaged by switching the filter cube (FESH 0899 for excitation, FELH 0900 for emission, Thorlabs) and using a mercury-vapor lamp (Olympus) for excitation.

### Electrical and optical stimulation-evoked dopamine imaging with near-infrared microscopy

For electrical stimulation experiments, a bipolar stimulation electrode was positioned in field of view within the dorsomedial striatum identified using a 4X objective (Olympus xFluor 4x/340). Using 60X objective, the stimulation electrode was brought into contact with top surface of the brain slice and an imaging field of view was chosen at a nominal distance of 150 μm from the stimulation electrode within the dorsomedial striatum. All stimulation experiments were recorded at video frame rates of 9 frames per second (nominal) and single pulse electrical stimulations were applied after 200 frames of baseline were acquired. Each video acquisition lasted 600 frames. Stimulation amplitudes were staggered and each stimulation amplitude was repeated three times within a field of view. Slices were allowed to recover for 5 minutes between each stimulation with the excitation laser path shuttered. For optogenetic stimulation, a fiber-coupled 473 nm blue laser (OptoEngine LLC DPSS) was positioned in close proximity to the brain slice using a micromanipulator. Expression of ChR2 was confirmed via visible fluorescence imaging and an imaging field of view was chosen in dorsomedial striatum with robust expression level. Stimulation pulses (5 pulses, 5 ms duration per pulse, delivered at 25 Hz, 1 mW/mm^2^) were delivered after acquiring 200 baseline frames and the video acquisition lasted 600 frames at nominal 9 frames per second. Drugs were bath applied to the imaging chamber through ACSF perfusion. ACSF with 10 μM of nomifensine or 1 μM of each DRD drug was used. When the effect of a drug needed to be evaluated, stimulation/imaging experiments were carried out with drug-free ACSF in an imaging field of view to collect drug-free data. Normal ACSF was then switched to ACSF prepared with the drug of interest and applied for 10 minutes before stimulation/imaging experiments resumed.

### Viral transfection of mice for optogenetic stimulation

Adult male and female mice (>P60) were used for all surgeries. Bilateral viral injections were performed using previously described procedures ^47^ at the following stereotaxic coordinates: dorsomedial prefrontal cortex (dmPFC): 1.94 mm from Bregma, 0.34 mm lateral from midline, and 0.70 mm vertical from cortical surface; substantia nigra pars compacta (SNc): −3.08 mm from Bregma, 1.25 mm lateral from midline, and 4.0 mm vertical from cortical surface. For glutamatergic corticostriatal axon stimulation experiments, mice were injected with 0.5 μL of CAG-ChR2-EYFP virus bilaterally into dmPFC. For nigrostriatal dopaminergic axon stimulation experiments, DAT-Cre mice were injected with 0.5 μL DIO-ChR2-EYFP virus bilaterally. For all optogenetic experiments, we waited at least three weeks from viral injection to experimental stimulation to allow for sufficient ChR2 gene expression.

To confirm that dopamine neurons were transfected with ChR2 in animals used for optogenetic dopamine stimulation experiments, we perfused DAT-Cre mice that had been injected into the SNc with Cre-dependent ChR2-EYFP virus with 4% paraformaldehyde in PBS and post-fixed brains overnight. Coronal sections that included the injection site (SNc) and imaging site (dorsal striatum) were cut at 50 μM and immunolabeled using antibody against tyrosine hydroxylase (TH) (rabbit anti-TH 1:1000, Millipore), the rate-limiting enzyme for catecholamine synthesis. Goat anti-rabbit Dylight 594 secondary antibody (1:1000, Invitrogen) was used to visualize TH. Image acquisition was performed on a Zeiss Axio ScanZ.1 using a 5x objective.

### Image processing and data analysis of nIRCat fluorescence response

Raw movie files were processed using a custom-built MATLAB program (available for download: https://github.com/jtdbod/Nanosensor-Brain-Imaging). Briefly, for each raw movie stack (600 frames), a per pixel *(F-F_0_)/F_0_* was calculated using the average intensity for the first 5% of frames as F_0_, and F represents the dynamic fluorescence intensity at each pixel. Regions of interest (ROIs) were identified by calculating a median filter convolution and then performing thresholding using Otsu’s method followed by a morphological dilation operation. ∆F/F traces were then calculated for each generated ROI. ROI sizes were computed using the measured pixel area and approximating each as a circle to calculate an equivalent radius.

To compare responses across stimulation amplitudes and bath application of nomifensine, mean results were obtained as follows: First, all identified ROIs from a field of imaging were averaged. Mean traces were further averaged over different fields of view within the same slice and across slices (1-2 field of view per slice, 1-2 slices per animal) and then averaged over experimental animals. Decay time constants (τ) were computed by fitting ∆F/F time traces to a first order decay process. Latency to peak were computed as *t_peak_* - *t_stim_* where *t_peak_* is the time at which peak signal is attained and *t_stim_* is time of stimulation. All statistical tests of significance (p-values) were computed and reported from unpaired, two-tailed t-test.

## Acknowledgements

We acknowledge support of an NIH NIDA CEBRA award # R21DA044010 (to L.W.), a Burroughs Wellcome Fund Career Award at the Scientific Interface (CASI) (to M.P.L.), the Simons Foundation (to M.P.L.), a Stanley Fahn PDF Junior Faculty Grant with Award # PF-JFA-1760 (to M.P.L.), a Beckman Foundation Young Investigator Award (to M.P.L.), and a DARPA Young Investigator Award (to M.P.L.). M.P.L. is a Chan Zuckerberg Biohub investigator. A.G.B. is supported by NSF Graduate Research Fellowship and an NIH DSPAN F99/K00 grant from NINDS. The authors wish to thank Marla Feller, Helen Bateup, David Sulzer, and David Schaffer for helpful discussions and comments on the manuscript and to Nicholas Ouassil for manuscript feedback.

## Data Availability Statement

All data as well as data processing software are available upon reasonable request to the corresponding author, and custom processing software is available on GitHub.

**Supplementary Information**

## References

1. Anderson, B. A. et al. The role of dopamine in value-based attentional orienting. Curr. Biol. (2016). doi:10.1016/j.cub.2015.12.062

2. Solanto, M. V. Dopamine dysfunction in AD/HD: Integrating clinical and basic neuroscience research. in Behavioural Brain Research (2002). doi:10.1016/S0166-4328(01)00431-4

3. Cohen, J. & Uchida, N. Neuron-type specific signals for reward and punishment in the ventral tegmental area. Nature (2012). doi:10.1038/nature10754.Neuron-type

4. Eshel, N., Tian, J., Bukwich, M. & Uchida, N. Dopamine neurons share common response function for reward prediction error. Nat. Neurosci. (2016). doi:10.1038/nn.4239

5. Berke, J. D. What does dopamine mean? Nature Neuroscience (2018). doi:10.1038/s41593-018-0152-y

6. Steinberg, E. E. et al. A causal link between prediction errors, dopamine neurons and learning. Nat. Neurosci. (2013). doi:10.1038/nn.3413

7. Hamid, A. A. et al. Mesolimbic dopamine signals the value of work. Nat. Neurosci. (2015). doi:10.1038/nn.4173

8. Salamone, J. D. & Correa, M. The Mysterious Motivational Functions of Mesolimbic Dopamine. Neuron (2012). doi:10.1016/j.neuron.2012.10.021

9. Dudman, J. T. & Krakauer, J. W. The basal ganglia: From motor commands to the control of vigor. Current Opinion in Neurobiology (2016). doi:10.1016/j.conb.2016.02.005

10. Lotharius, J. & Brundin, P. Pathogenesis of parkinson’s disease: Dopamine, vesicles and α-synuclein. Nat. Rev. Neurosci. (2002). doi:10.1038/nrn983

11. Weinstein, J. J. et al. Pathway-Specific Dopamine Abnormalities in Schizophrenia. Biological Psychiatry (2017). doi:10.1016/j.biopsych.2016.03.2104

12. Volkow, N. D., Fowler, J. S., Wang, G.-J. & Swanson, J. M. Dopamine in drug abuse and addiction: results from imaging studies and treatment implications. Mol. Psychiatry (2004). doi:10.1038/sj.mp.4001507

13. Clements, J., Lester, R., Tong, G., Jahr, C. & Westbrook, G. The time course of glutamate in the synaptic cleft. Science (80-.). (1992). doi:10.1126/science.1359647

14. Greengard, P. The neurobiology of slow synaptic transmission. Science (80-.). (2001). doi:10.1126/science.294.5544.1024

15. Agnati, L. F., Zoli, M., Strömberg, I. & Fuxe, K. Intercellular communication in the brain: Wiring versus volume transmission. Neuroscience (1995). doi:10.1016/0306-4522(95)00308-6

16. Zoli, M. et al. The emergence of the volume transmission concept. in Brain Research Reviews (1998). doi:10.1016/S0165-0173(97)00048-9

17. Rice, M. E. & Cragg, S. J. Dopamine spillover after quantal release: Rethinking dopamine transmission in the nigrostriatal pathway. Brain Research Reviews (2008). doi:10.1016/j.brainresrev.2008.02.004

18. Cragg, S. J. & Rice, M. E. DAncing past the DAT at a DA synapse. Trends in Neurosciences (2004). doi:10.1016/j.tins.2004.03.011

19. Dreyer, J. K., Herrik, K. F., Berg, R. W. & Hounsgaard, J. D. Influence of Phasic and Tonic Dopamine Release on Receptor Activation. J. Neurosci. (2010). doi:10.1523/JNEUROSCI.1894-10.2010

20. Marder, E. Neuromodulation of Neuronal Circuits: Back to the Future. Neuron (2012). doi:10.1016/j.neuron.2012.09.010

21. Hong, G. et al. Through-skull fluorescence imaging of the brain in a new near-infrared window. Nat. Photonics (2014). doi:10.1038/nphoton.2014.166

22. Zhang, J. et al. Molecular recognition using corona phase complexes made of synthetic polymers adsorbed on carbon nanotubes. Nat. Nanotechnol. (2013). doi:10.1038/nnano.2013.236

23. Kruss, S. et al. Neurotransmitter detection using corona phase molecular recognition on fluorescent single-walled carbon nanotube sensors. J. Am. Chem. Soc. (2014). doi:10.1021/ja410433b

24. Beyene, A. G., McFarlane, I. R., Pinals, R. L. & Landry, M. P. Stochastic Simulation of Dopamine Neuromodulation for Implementation of Fluorescent Neurochemical Probes in the Striatal Extracellular Space. ACS Chem. Neurosci. (2017). doi:10.1021/acschemneuro.7b00193

25. Berridge, C. W. & Waterhouse, B. D. The locus coeruleus-noradrenergic system: Modulation of behavioral state and state-dependent cognitive processes. Brain Research Reviews (2003). doi:10.1016/S0165-0173(03)00143-7

26. Gerfen, C. R. Synaptic organization of the striatum. J. Electron Microsc. Tech. (1988). doi:10.1002/jemt.1060100305

27. Tritsch, N. X. & Sabatini, B. L. Dopaminergic Modulation of Synaptic Transmission in Cortex and Striatum. Neuron (2012). doi:10.1016/j.neuron.2012.09.023

28. Piekarski, D. J., Boivin, J. R. & Wilbrecht, L. Ovarian Hormones Organize the Maturation of Inhibitory Neurotransmission in the Frontal Cortex at Puberty Onset in Female Mice. Curr. Biol. (2017). doi:10.1016/j.cub.2017.05.027

29. Godin, A. G. et al. Single-nanotube tracking reveals the nanoscale organization of the extracellular space in the live brain. Nat. Nanotechnol. (2017). doi:10.1038/nnano.2016.248

30. Robinson, D. L., Venton, B. J., Heien, M. L. A. V & Wightman, R. M. Detecting subsecond dopamine release with fast-scan cyclic voltammetry in vivo. in Clinical Chemistry (2003). doi:10.1373/49.10.1763

31. Heien, M. L. A. V., Johnson, M. A. & Wightman, R. M. Resolving neurotransmitters detected by fast-scan cyclic voltammetry. Anal. Chem. (2004). doi:10.1021/ac0491509

32. Garris, P. A. & Wightman, R. M. Regional Differences in Dopamine Release, Uptake, and Diffusion Measured by Fast-Scan Cyclic Voltammetry. Neuromethods: Voltammetric Methods in Brain Systems (1995). doi:10.1385/0-89603-312-0:179

33. Sun, F. et al. A Genetically Encoded Fluorescent Sensor Enables Rapid and Specific Detection of Dopamine in Flies, Fish, and Mice. Cell (2018). doi:10.1016/j.cell.2018.06.042

34. Patriarchi, T. et al. Ultrafast neuronal imaging of dopamine dynamics with designed genetically encoded sensors. Science (80-.). (2018). doi:10.1126/science.aat4422

35. Pereira, D. B. et al. Fluorescent false neurotransmitter reveals functionally silent dopamine vesicle clusters in the striatum. Nat. Neurosci. (2016). doi:10.1038/nn.4252

36. Liu, C., Kershberg, L., Wang, J., Schneeberger, S. & Kaeser, P. S. Dopamine Secretion Is Mediated by Sparse Active Zone-like Release Sites. Cell 172, 706–718.e15 (2018).

37. Mohebi, A. et al. Forebrain dopamine value signals arise independently from midbrain dopamine cell firing. bioRxiv (2018). doi:10.1101/334060

38. Heller, D. a et al. Optical detection of DNA conformational polymorphism on single-walled carbon nanotubes. Science (2006). doi:10.1126/science.1120792

39. Cognet, L. et al. Stepwise quenching of exciton fluorescence in carbon nanotubes by single-molecule reactions. Science (80-.). (2007). doi:10.1126/science.1141316

40. Barone, P. W., Baik, S., Heller, D. A. & Strano, M. S. Near-infrared optical sensors based on single-walled carbon nanotubes. Nat. Mater. (2005). doi:10.1038/nmat1276

41. Roxbury, D. et al. Hyperspectral Microscopy of Near-Infrared Fluorescence Enables 17-Chirality Carbon Nanotube Imaging. Sci. Rep. (2015). doi:10.1038/srep14167

42. Giraldo, J. P. et al. A Ratiometric Sensor Using Single Chirality Near-Infrared Fluorescent Carbon Nanotubes: Application to In Vivo Monitoring. Small (2015). doi:10.1002/smll.201403276

43. Bonis-O’Donnell, J. T. D. et al. Dual Near-Infrared Two-Photon Microscopy for Deep-Tissue Dopamine Nanosensor Imaging. Adv. Funct. Mater. (2017). doi:10.1002/adfm.201702112

44. Ding, J. B., Takasaki, K. T. & Sabatini, B. L. Supraresolution Imaging in Brain Slices using Stimulated-Emission Depletion Two-Photon Laser Scanning Microscopy. Neuron (2009). doi:10.1016/j.neuron.2009.07.011

45. Wang, F., Dukovic, G., Brus, L. E. & Heinz, T. F. The optical resonances in carbon nanotubes arise from excitons. Science (80-.). (2005). doi:10.1126/science.1110265

46. Edelstein, A. D. et al. Advanced methods of microscope control using μManager software. J. Biol. Methods (2014). doi:10.14440/jbm.2014.36

47. Vandenberg, A., Piekarski, D. J., Caporale, N., Munoz-Cuevas, F. J. & Wilbrecht, L. Adolescent maturation of inhibitory inputs onto cingulate cortex neurons is cell-type specific and TrkB dependent. Front. Neural Circuits (2015). doi:10.3389/fncir.2015.00005

